# Subcellular Level Spatial Transcriptomics with PHOTON

**DOI:** 10.1101/2024.09.10.612328

**Authors:** Shreya Rajachandran, Qianlan Xu, Qiqi Cao, Xin Zhang, Fei Chen, Sarah M. Mangiameli, Haiqi Chen

## Abstract

The subcellular localization of RNA is closely linked to its function. Many RNA species are partitioned into organelles and other subcellular compartments for storage, processing, translation, or degradation. Thus, capturing the subcellular spatial distribution of RNA would directly contribute to the understanding of RNA functions and regulation. Here, we present PHOTON (Photoselection of Transcriptome over Nanoscale), a method which combines high resolution imaging with high throughput sequencing to achieve spatial transcriptome profiling at subcellular resolution. We demonstrate PHOTON as a versatile tool to accurately capture the transcriptome of target cell types *in situ* at the tissue level such as granulosa cells in the ovary, as well as RNA content within subcellular compartments such as the nucleolus and the stress granule. Using PHOTON, we also reveal the functional role of m^6^A modification on mRNA partitioning into stress granules. These results collectively demonstrate that PHOTON is a flexible and generalizable platform for understanding subcellular molecular dynamics through the transcriptomic lens.

## Introduction

RNA distribution within a cell is intimately linked to cell functions [1, 2]. The enrichment of selective RNA species in subcellular compartments is a phenomenon observed across species. These RNA-containing compartments partition cellular content, create cellular asymmetries, enhance biological reactions, and promote molecular interactions required for cellular functions and cell fate decisions [3-5]. For instance, selective mRNAs have been shown to be recruited into the stress granules (SGs) for translation suppression during cellular stress [6, 7]. In the nucleus, nucleolus has been demonstrated to be the site for ribosomal RNA (rRNA) synthesis [8] and inflammatory RNA decay during infection [9]. Given the significant roles of RNA distribution in cell functions, there is a need for tools that can specifically capture RNA species within subcellular compartments.

While many methods have been developed to study RNA distribution within cells, only a few have been applied on a transcriptome-wide scale. Furthermore, current transcriptomics methods suffer from various limitations. For example, a conventional approach to profile RNA content within subcellular compartments is to biochemically (e.g., protein-RNA crosslinking followed by immunoprecipitation) and/or mechanically (e.g., density gradient followed by centrifugation) purify these compartments and sequence their RNA content [10-12]. However, this approach requires millions of cells as input, which makes it challenging to apply to rare cell types. And more importantly, not all subcellular compartments can be purified, especially for those membrane-less, transient biomolecular condensates. Even for compartments that can be isolated, such as the SGs, current protocols fail to remove contaminant or prevent content loss during the isolation process.

Imaging-based spatial transcriptomics methods such as MERFISH [13] and seqFISH [14] can visualize the distribution of thousands of mRNAs within individual cells. The drawbacks of these approaches, however, are the need for designing and synthesizing a large pool of probe sets targeting RNAs of interest, the requirement for specialized technical expertise and instrumentation, and sophisticated image processing and data analysis.

Emerging proximity labeling-based RNA profiling methods such as APEX-seq are powerful tools to capture RNA transcripts at a high spatiotemporal resolution [15, 16]. However, they often require time-consuming genome engineering in which a labeling enzyme (e.g., a biotin ligase) is genetically fused to a protein enriched in a type of compartment. This makes it difficult to scale to various types of subcellular compartments. And the specificity of these methods is dictated by the specificity of the protein the labeling enzyme is fused to.

Thus, there remains a need for new tools that 1) can capture the spatial localization of thousands of endogenous RNA species within various types of subcellular compartments; 2) require significantly less input than exisiting methods; and 3) do not need genetic manipulations. Here, we present PHOTON (<u>Ph</u>otoselection <u>o</u>f <u>T</u>ranscriptome <u>o</u>ver <u>N</u>anoscale), a method which combines high resolution imaging with high throughput sequencing to achieve spatial transcriptome profiling at subcellular resolution. We demonstrate PHOTON as a versatile tool to capture the transcriptome of target cell types *in situ* at the tissue level as well as RNA content within subcellular compartments. Using PHOTON, we also reveal the functional role of m^6^A modification on mRNA partitioning into SGs. These examples illustrate the versatility of PHOTON and its ability to reveal new biological insights.

## Results

### The PHOTON Workflow

To develop the methodology, we drew from our previous work in sequencing DNA of fixed samples at sub-micron (∼300 nm) resolution using imaging-guided, laser targeted photo-selection [17]. Based on the same imaging setup, PHOTON can be broken down into four major steps: 1) construction of a photocaged cDNA library *in situ*; 2) selective uncaging of the cDNA molecules using targeted illumination with near-ultraviolet (UV) laser light; 3) sample digestion and PCR handle ligation to uncaged cDNA molecules; and 4) library preparation followed by sequencing on an Illumina platform (**Figure 1A**).

**Figure 1.**
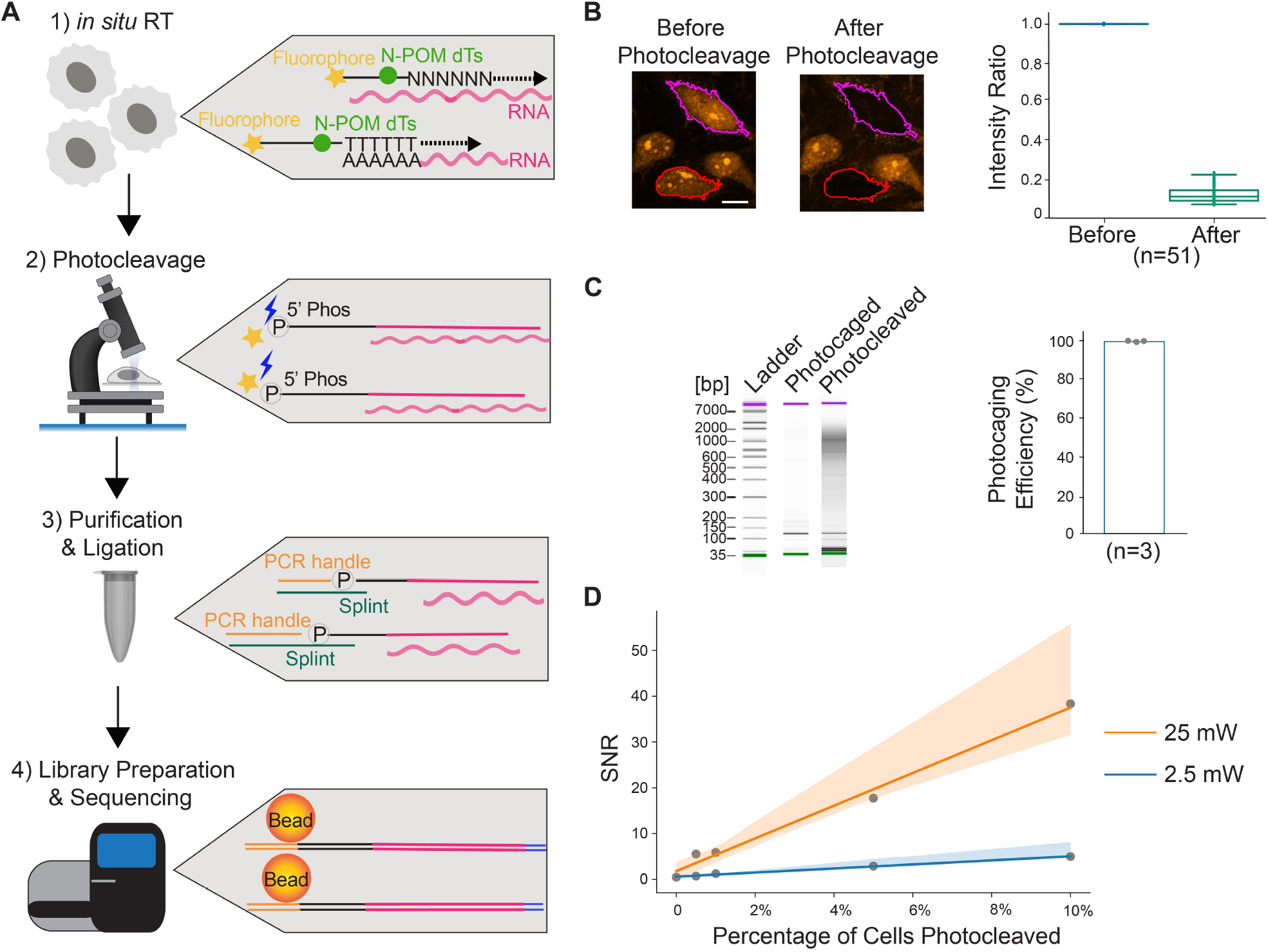
PHOTON enables spatially resolved transcriptome profiling. **(A)**Schematic diagram illustrating the PHOTON method. First, *in situ* RT is performed using photocleavable RT primers. Second, targeted illumination is performed within specific ROIs identified by fluorescence imaging. Exposure of the cDNA molecules to near-UV light modifies the N-POM-caged dTs and breaks the photocleavable linkers on the RT primers, releasing the fluorescent labels and revealing the phosphate groups. Third, nucleic acids are purified, and PCR handles are ligated to previously photocleaved cDNA molecules through splint oligos. Finally, successfully ligated cDNA molecules are pulled down using streptavidin beads and a template switching step is performed with TSOs. Following PCR amplification, sequencing libraries are generated and read by next-generation sequencing. **(B)**Photocleavage of cDNA molecules in HeLa cells. The color-outlined cells were exposed to 25 mW 405 nm laser light which cleaved the fluorophores from the RT primers and resulted in an 86.9% (±4.3% s.d.) intensity decrease (n = 51 cells, 1 experiment). Scale bar, 5 µm. **(C)**Measuring the efficiency of the PHOTON photocaging mechanism. Left: tapestation gel image of a representative experiment; Right: bar graph showing the photocaging efficiency of PHOTON (n = 3 experiments). On average, 99.1 ± 0.26% of the photocaged cDNA molecules could not be amplified without near-UV light exposure. **(D)**SNR of PHOTON as a function of the fraction of cells selected under two different laser powers.

First, to construct a photocaged *in situ* cDNA library, we use custom reverse transcription (RT) primers. These primers contain, at the 3’ end, either a poly dT sequence or a random hexamer to hybridize to both mRNA and other non-poly adenylated RNA species. The 5’ end of the RT primer is linked to a fluorophore using a photocleavable linker. In addition, four 6-nitropiperonyloxymethyl (N-POM)-modified dTs are also incorporated into the 5’ end of the primers (**Figure S1A; Table S1**).

We next visualize the sample by microscopy and use fluorescent stains of a subcellular compartment to guide the identification of regions of interest (ROIs). Automated in-line image segmentation allows scalable assays of thousands of individual ROIs localized throughout the sample. Selective illumination of the ROIs using a 405 nm laser line cleaves the fluorophores from the RT primers via photocleavage of the linker, revealing a 5’ phosphate group. In the meantime, the 405 nm laser line also uncages the N-POM dTs within the primers, restoring the base paring functions of the dTs (**Figure 1A**, Step 2). Thus, the photocleavable linker and the N-POM dTs serve as dual photocaging mechanisms to prevent unwanted uncaging of cDNA molecules.

Following photocleavage, cells are lysed, and nucleic acids are purified. A PCR handle containing a priming site for library amplification is then ligated to the 5’ end of the RT primer through a splint oligo. This ligation step only happens to the cDNA molecule where the 5’ phosphate group was previously revealed and the N-POM dTs were uncaged through photocleavage (**Figure 1A**, Step 3). Photocaged cDNA molecules do not have available 5’ phosphates and the intact N-POM dTs prevent the base pairing of the splint oligos to the RT primers. Therefore, the photocaged cDNA molecules cannot pass the ligation stage.

Successfully ligated cDNA molecules can be pulled down by streptavidin beads due to the presence of biotin molecules on the PCR handle. These purified cDNA molecules are subject to template switching using template switching oligos (TSOs) to add a second PCR handle. Following PCR amplification, the PCR products are subject to Illumina Nextera XT sequencing library preparation workflow and the resulting libraries are depleted of rRNA sequences using DASH [18]. The final products are sequenced on an Illumina platform (**Figure 1A**, Step 4).

### The Feasibility of PHOTON

To test the feasibility of PHOTON, we first characterized PHOTON libraries in cultured HeLa cells. To do this, we treated fixed and permeabilized HeLa cells with the custom photocaged RT primers and performed *in situ* RT. cDNA products were visualized through the fluorophore on the RT primers (**Figure 1B**, left image). When a subset of cells was exposed to a focused 405 nm laser light (**Figure 1B**, images), we observed an accompanying 86.9% (±4.3% standard deviation (s.d.), N = 51 cells) decrease in fluorescence intensity within the exposed cells (**Figure 1B**, plot) due to the photocleavage and subsequent diffusion of the fluorophore. In contrast, no decrease in fluorescence intensity was observed in non-targeted cells (**Figure S1B**).

To confirm that PHOTON libraries were generated from cellular RNA, we subjected two groups of cells (∼5,000 cells in each group) to the PHOTON workflow except that the reverse transcriptase was omitted in one group of cells (i.e., no RT). As expected, no PHOTON library was generated from the no RT group (**Figure S1C**).

To assess the baseline efficacy of the photocaging mechanism of the custom RT primers, we compared the size of PHOTON libraries generated with and without photocleaving the RT primers. Briefly, following *in situ* RT using the RT primers, cDNA products were isolated from the cells and split into halves. One half was exposed to the 405 nm laser light to photocleave the RT primers while the other half was protected from light. The two halves were then subject to the rest of the PHOTON workflow. As expected, the photocleaved half resulted in a PHOTON library while the photocaged half did not (**Figure 1C**, left). Quantification of the two halves using qPCR showed that the photocaging mechanism is 99.1 ± 0.26% (n=3 experiments) effective in preventing unwanted photocleavage of cDNA molecules (**Figure 1C**, right).

We next quantified the signal-to-noise ratio (SNR) of PHOTON. Given the design of PHOTON, we expect that the noise mainly stems from unwanted photocleavage of cDNA molecules due to, for instance, accidental exposure of the photocaged cDNA molecules to ambient light during sample processing or defective RT primer oligos during oligo synthesis. We also expect that the amount of signals scale with the level of laser power (stronger laser power means that more cDNA molecules can be photocleaved), as well as the number of cells photocleaved (which is proportional to the area of ROIs). Therefore, we calculated the relationship between the SNR and the laser power as well as the fraction of cells photocleaved *in situ*. To do this, we photocleaved various proportions of ∼5,000 HeLa cells using two different levels of laser power and measured the SNR under each condition. We found that under the 25 mW laser light, the SNR was approximately 357 times the fraction of cells photocleaved; and under 2.5 mW laser light, the SNR was 57 times the fraction of cells photocleaved. Notably, the signal of PHOTON was still above the noise (SNR>1.2) even though only 0.5% of the cells (∼25 HeLa cells) were photocleaved at a low laser power of 2.5 mW (**Figure 1D**).

Together, these results suggest that PHOTON is highly sensitive in capturing RNA information *in situ*.

### PHOTON Reproduces Transcriptome Data in the Mouse Ovary

After demonstrating the feasibility of PHOTON, we first set out to validate its capability to capture cellular transcriptome at the tissue level. We focused on granulosa cells (GCs; a cell type supporting oocyte development) in the ovarian follicles because GCs can be easily identified through their spatial locations and tissue morphology (**Figure 2A** & **Figure S2A**). We generated two sets of PHOTON libraries by each photocleaving approximately 600 GCs from follicles at various developmental stages in a mouse ovarian tissue section. The PHOTON data were highly reproducible, as demonstrated by the high correlation between two replicates (**Figure S2B**). Transcriptome analysis showed that known GC markers such as *Inha, Nr5a2*, and *Serpine2* were highly expressed in the PHOTON datasets while markers of other cell types in the ovary were not (**Figure 2B**).

**Figure 2.**
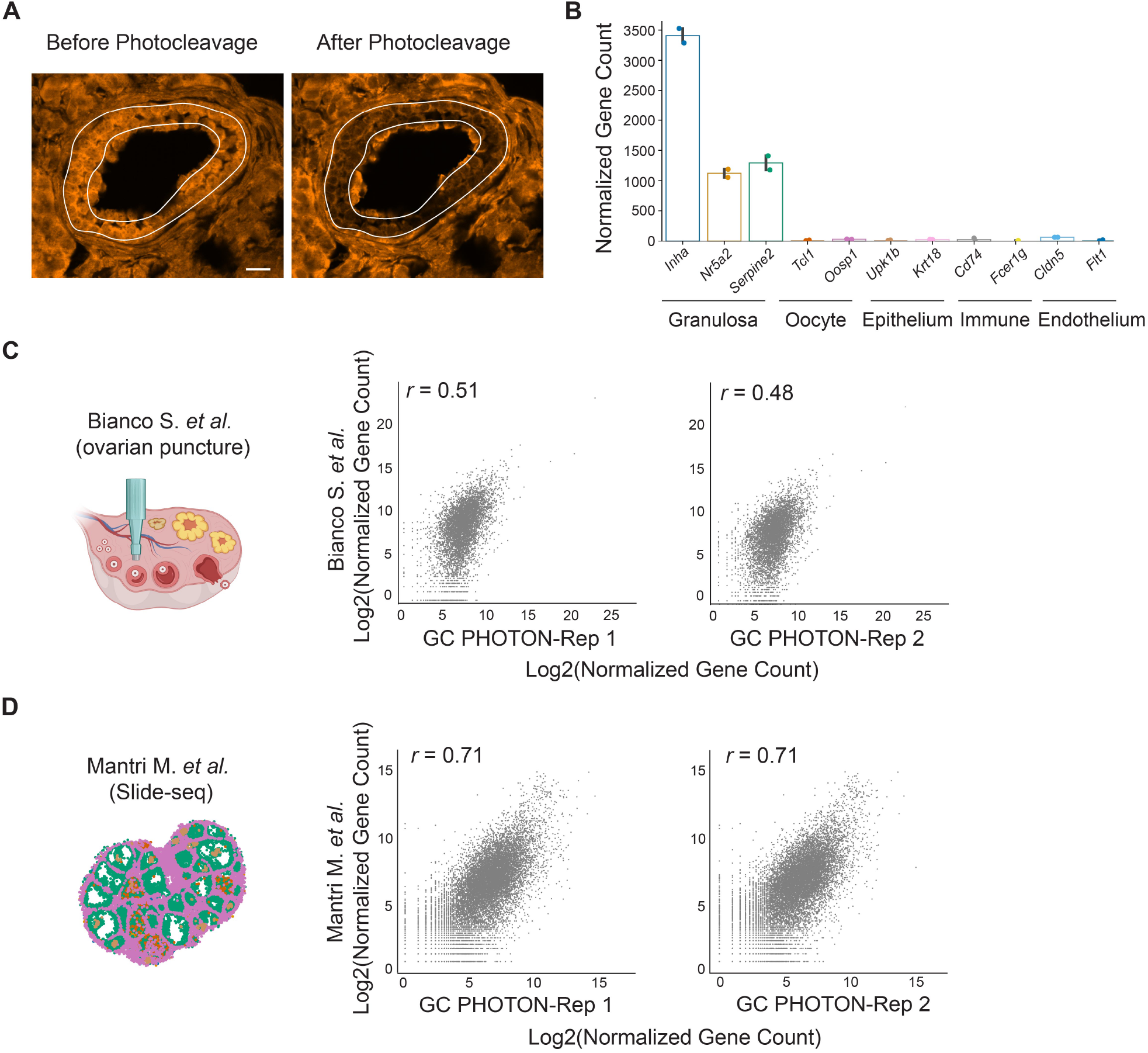
PHOTON captures spatial transcriptomics data of granulosa cells in the mouse ovary. **(A)**Morphologically guided targeting of GCs in the adult mouse ovarian tissue slices. A secondary follicle was visualized by the fluorescence of the photocaged cDNA molecules. Scale bar, 15 µm. **(B)**Expression levels of ovarian cell type marker genes revealed by PHOTON. **(C)**Comparison between the mouse GC transcriptome generated by the follicle isolation method and that generated by PHOTON. **(D)**Comparison between the mouse GC transcriptome generated by Slide-seqV2 and that generated by PHOTON.

To further benchmark the GC transcriptome data generated using PHOTON, we compared them to a publicly available GC RNA-seq dataset. This public dataset was generated by mechanically dissecting out individual follicles from mouse ovarian tissues using punctures and then isolating GCs from follicles [19]. Both PHOTON replicates moderately correlated with the public RNA-seq dataset (**Figure 2C**). We reasoned that this was because the mechanical method of isolating GCs could not fully tease out other ovarian cell types and therefore the reference library may contain contaminates. To this end, we extracted the GC transcriptome from another publicly available dataset in which mouse ovarian tissue sections were subject to unbiased spatial transcriptome profiling at a near single cell resolution using Slide-seqV2 [20]. This approach does not require GC isolation. Indeed, a much higher correlation coefficient was observed between the PHOTON replicates and the Slide-seqV2 data (**Figure 2D**).

Together, these data suggest that PHOTON can accurately capture the transcriptome information of target cell types within a tissue in a spatially resolved manner.

### PHOTON Assays Transcriptome in Subcellular Compartments

Following the validation of PHOTON at the tissue level, we next sought to validate its capability to capture RNA information at the subcellular level. We first turned to nucleolus – a well characterized nuclear compartment with an essential role in ribosome biogenesis. We developed a computational pipeline to automatically identify and photocleave cDNA molecules in the nucleolus of fixed HeLa cells (**Figure 3A**). The nucleolus PHOTON data was highly reproducible as demonstrated by the high correlation between the two replicates (**Figure S3A**). In parallel, we also used PHOTON to generate paired whole cell transcriptome data. This allowed us to identify RNA species that were specifically enriched in the nucleolus. The nucleolus is known to be rich in noncoding RNAs, mostly the rRNAs and small nucleolar RNAs (snoRNAs). Indeed, the majority of top-enriched RNAs identified by PHOTON were snoRNAs (because DASH, an rRNA depletion protocol, was included in the PHOTON workflow, rRNA was not detected in the PHOTON datasets) (**Figure 3B**). Of interest, PHOTON also identified long noncoding RNA *RMRP* to be associated with the nucleolus (**Figure 3B**), consistent with a previous finding [21].

**Figure 3.**
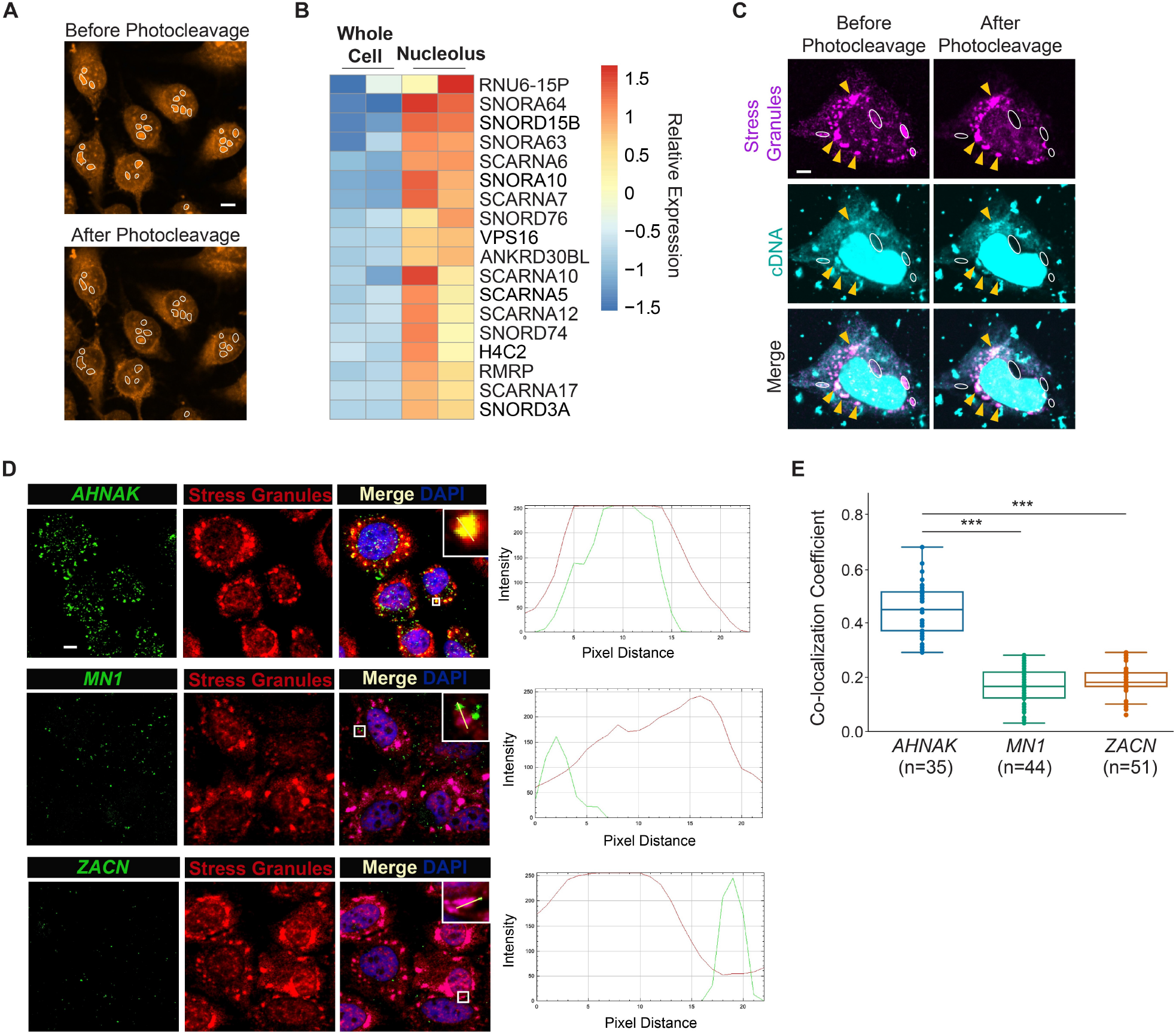
PHOTON captures transcriptome information within subcellular compartments. **(A)**Representative images of photocleaving cDNA molecules within nucleoli (white outlines). Scale bar, 2 µm. **(B)**Heatmap showing the top-enriched RNA species in the nucleolus identified using the PHOTON data. **(C)**Representative images of photocleaving cDNA molecules (cyan) in SGs (magenta). SGs with white dashed outlines were targeted for photocleavage, while others such as those with yellow arrow heads were not. Scale bar, 1 µm. **(D)**HCR-FISH images showing the spatial distribution of *AHNAK, MN1*, and *ZACN* mRNAs (green) in relation to SGs (red). Line graphs on the right show the intensity profiles of both the mRNA signal (green line) and the SG signal (red line) along the yellow line in the inserts. Scale bar, 2 µm. **(E)**Co-localization coefficient of the mRNA and SG signals for *AHNAK, MN1*, and *ZACN*. ^***^, *p* <0.001. One-way ANOVA followed by Tukey test.

To further showcase the versatility of PHOTON in characterizing subcellular compartments, we next focused on SGs. SGs are cytoplasmic mRNA-protein assemblies that are important in the cellular stress response and may contribute to degenerative diseases [22]. We applied PHOTON to capture the RNA content of SGs in HeLa cells induced by sodium arsenite treatment (**Figure 3C**). The SG PHOTON data was highly reproducible as demonstrated by the high correlation between three replicates (**Figure S3B**). By pairing the SG PHOTON data with the whole cell PHOTON data, we identified transcripts that were differentially enriched and depleted in the SGs (|Log2 fold change|>1; adjusted p value <0.05). When comparing these data with a publicly available SG dataset generated by the conventional purification method [10], we found a significant overlap (*p* < 0.001, hypergeometric test) between the SG-enriched/SG-depleted transcripts from the PHOTON data and those from the public data (742 overlapped SG-enriched genes out of 1616 genes; 864 overlapped SG-depleted genes out of 1780 genes).

We next performed hybridization chain reaction-based fluorescence in situ hybridization (HCR-FISH) to verify the subcellular localization of some of the identified transcripts. For example, *AHNAK* mRNA was found to be enriched in SGs in both the PHOTON data and the public dataset. This observation was confirmed by HCR-FISH imaging of the *AHNAK* transcripts which showed a significant overlap between the *AHNAK* mRNA signals and SG signals (**Figure 3D**, upper panel; **Figure 3E**). Of interest, there were also discrepancies between the PHOTON data and the public dataset. For instance, both *MN1* and *ZACN* mRNAs were reported to be enriched in SGs in the public dataset. However, our PHOTON data indicated no enrichment of these two mRNA in the SGs. Indeed, HCR-FISH showed that neither *MN1* nor *ZACN* mRNA signals significantly overlapped with the SG signals (**Figure 3D**, middle and lower panel; **Figure 3E**). This discrepancy between the PHOTON data and the public dataset may be due to the low specificity of the conventional SG purification method as contaminations might have occurred during the SG isolation procedure.

Together, these results suggest that PHOTON can accurately assay the RNA content of subcellular compartments.

### PHOTON Provides Insights into the Mechanism of mRNA Recruitment into SGs

Finally, we leveraged PHOTON to investigate the regulatory mechanisms governing RNA subcellular distributions.

Previous studies have demonstrated that during cellular stress, mRNAs with longer transcript length are preferentially enriched in SGs than shorter ones [10, 11]. Analysis of our SG PHOTON data from HeLa cells confirmed this observation (**Figure 4C**, left), which further validated the ability of PHOTON to accurately capture RNA content in subcellular compartments. Despite this length bias, mRNAs of the same length can show different recruitment into SGs, suggesting the existence of sequence-specific factors that affect mRNA recruitment into SGs. One of such length-independent factors have been proposed to be the *N*^6^-methyladenosine (m^6^A) modifications on mRNAs as the binding of YTHDF proteins to m^6^A may facilitate the partitioning of the bond mRNAs into SGs [23, 24] (**Figure 4A**). However, disentangling the relative contribution of m^6^A from that of the transcript length in mediating mRNA enrichment in SGs is challenging as m^6^A often occurs within long internal exons which are often present in long transcripts [25]. Furthermore, a recent study suggests that m^6^A modifications have negligible effects on mRNA partitioning into SGs [26].

**Figure 4.**
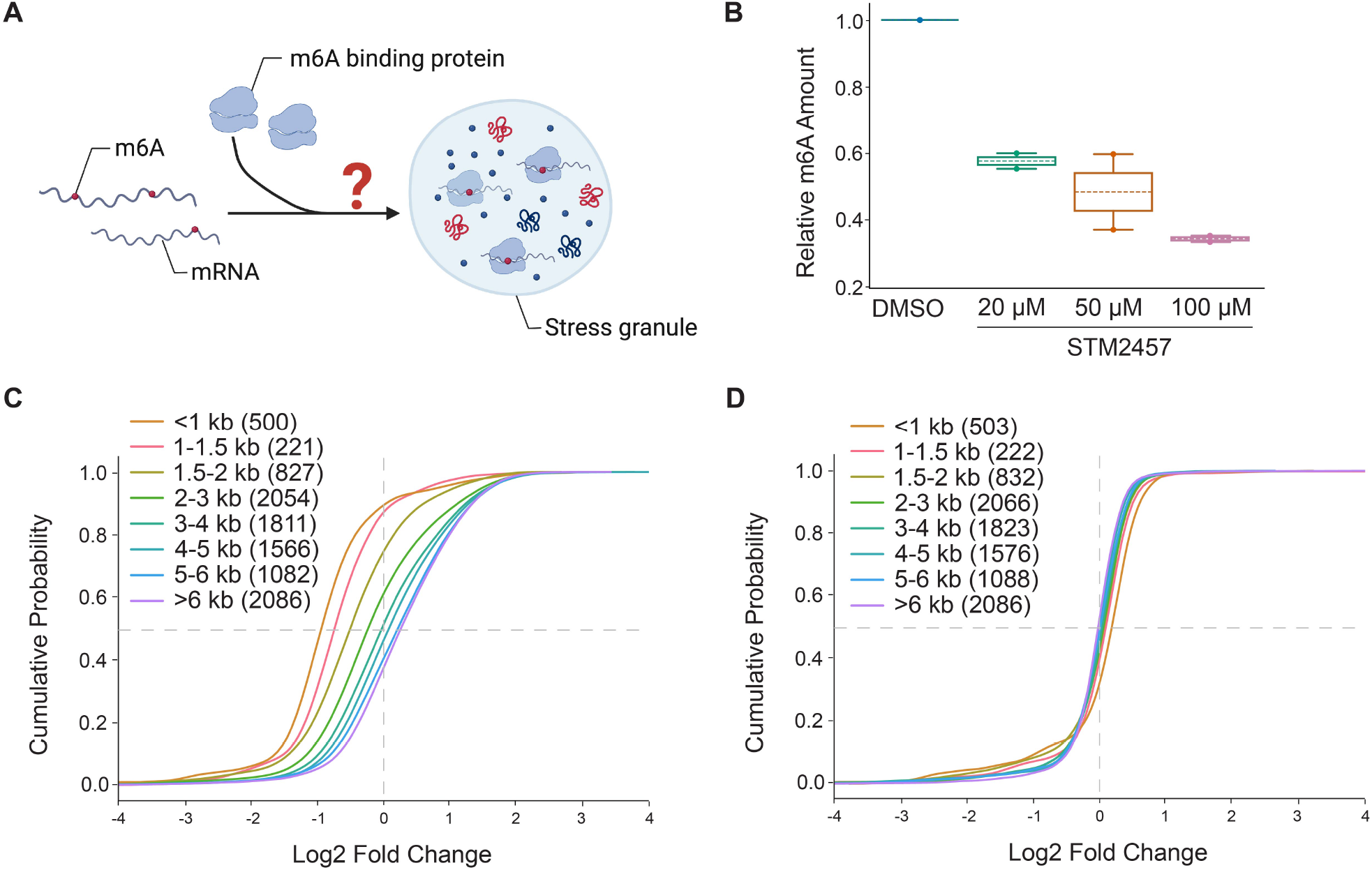
PHOTON reveals the functional role of m^6^A in mRNA recruitment into SGs. **(A)**m^6^A modifications may facilitate the partitioning of mRNAs into SGs through m^6^A-binding proteins. **(B)**Dose-dependent depletion of mRNA m^6^A modifications using METTL3 inhibitor STM2457. **(C)**Cumulative distribution plot of transcript abundance in the SGs of normal HeLa cells, binned based on the transcript length. Numbers in the parentheses indicate the number of transcripts in each bin. **(D)**Cumulative distribution plot of transcript abundance in SGs of STM2457 (100 µM)-treated HeLa cells, binned based on the transcript length. Numbers in the parentheses indicate the number of transcripts in each bin.

We therefore set out to use PHOTON to clarify the role of m^6^A modifications in the spatial distribution of mRNAs in relation to SGs. To this end, we first used a drug-inducible *Mettl3*-knockout mouse embryonic fibroblast (MEF) cell line (*Mettl3*KO). Previous work showed that 4-hydroxytamoxifen (4-OHT) treatment of this cell line for 7 days resulted in a loss of Mettl3 protein and subsequently a reduction of m^6^A on mRNAs [27]. We captured the RNA content of sodium arsenite-induced SGs in both wild type (WT) and *Mettl3*KO MEF cells using PHOTON while also capturing the corresponding total cellular RNA in parallel as input. We then focused on four genes with known quantities of m^6^A sites on their mRNAs. We quantified their fold changes in SG enrichment in *Mettl3*KO *vs*. WT cells using qPCR (n=3 replicates per gene). We found that there was a negative effect of m^6^A reduction on the enrichment of mRNA transcripts in SGs. This negative effect was increasingly pronounced in transcripts with more m^6^A modifications (**Figure S4**).

Next, we conducted a transcriptome-wide analysis to provide a comprehensive measurement of the role of m^6^A modifications in mRNA recruitment to SGs. To this end, we treated HeLa cells with a potent METTL3 inhibitor STM2457 to reduce the m^6^A level on mRNAs (**Figure 4B**) and induced SG formation using sodium arsenite. To distinguish the contribution of mRNA length and m^6^A in recruiting mRNA to SGs, mRNAs were grouped into bins based on the transcript length, and their fold enrichment in SGs was quantified using PHOTON. We found that, unlike the normal control HeLa cells shown in **Figure 4C**, the length-dependent mRNA enrichment in SG was lost in HeLa cells treated with STM2457 at 100 μM for 6 hours (**Figure 4D**) even though PHOTON captured a comparable number of transcripts in each transcript length bin between the control and the treatment group (n=3 experiments per group).

Together, these results suggest that the correlation between the transcript length and the SG enrichment is mediated, at least in part, by m^6^A modifications.

## Discussion

PHOTON reparents a new way of extracting subcellular transcriptomic information by assimilating both imaging and sequencing technologies. We demonstrated that PHOTON can be flexibly applied across ROIs that spanned order-of-magnitude size scales, from specific regions of the mouse ovarian tissue to the nucleolus and SGs within cells. In-line image segmentation enabled ROI generation based on an extensive range of spatial features and automated the targeted photocleavage process over large numbers of cells or features. At the tissue scale, PHOTON accurately captured the transcriptome of GCs within their native tissue microenvironment. At the subcellular scale, we used PHOTON to enable selective sequencing of the RNA content in the nucleolus and SGs and demonstrate the role of m^6^A modifications in the recruitment of mRNA into SGs. Taken together, these results show that PHOTON has the potential to uncover new connections between spatial and transcriptomic information at diverse length scales.

At the tissue level, we anticipate that PHOTON will be a useful alternative to existing spatial transcriptomics technology for fast, sensitive, and robust spatial annotation of cell types and states. Compared with PHOTON, existing spatial transcriptomics methods can be inaccessible due to the high cost. For example, array-based spatial barcoding methods powerfully profile large numbers of cells in tissue sections, but require the use of specialized microfluidics or custom arrays [28, 29]. Furthermore, because library generation is not targeted to a cell population of interest, sequencing costs can be high (especially for human samples), and deep sequencing is required to adequately profile uncommon cell types. In contrast, PHOTON only requires common reagents that can be directly ordered in bulk quantity from common vendors and a suitable imaging system that is available in most institutional microscopy cores. Furthermore, PHOTON reduces sequencing costs by targeting library generation to the cellular population of interest, which decreases the number of sequencing reads needed to adequately profile the sample.

At the subcellular level, PHOTON adds to the arsenal of RNA localization methods while offering unique advantages. First, PHOTON can be used to analyze ‘‘unpurifiable’’ structures such as the membrane-less biomolecular condensates that are challenging to access via conventional purification-based approaches. Second, PHOTON provides sequence information for both coding and non-coding RNA transcripts. If combined with long-read sequencing platforms, PHOTON can be readily adapted to allow transcript isoforms with distinct localization to be distinguished. Third, no genetic manipulation or antibody-based pulled down is needed in the PHOTON workflow, making it easy to scale while still highly specific. Finally, the high spatial resolution of the laser light employed in PHOTON allows the selective sequencing of ROIs with a spatial resolution down to the diffraction limit (200-300 nm).

Since many subcellular compartments directly contribute to the regulation of RNAs, PHOTON offers opportunities for mechanistic discoveries of this process. For instance, SGs have been described as a triage for mRNAs during cellular stress where they either store translationally silent mRNA, transfer mRNA transcripts to processing bodies where they will be degraded, or transfer mRNA back into polysomes for translation [30]. Thus, understanding which mRNAs are preferentially recruited to SGs during cellular stress and the underlying recruitment mechanisms would inform us on how cells cope with stress under physiological and pathological conditions. Using PHOTON, we confirmed a previous observation that long mRNAs tend to localize to SGs. Furthermore, we showed that this length-dependent enrichment of mRNAs in SGs was mediated, at least partially, by m^6^A as long mRNAs failed to show enrichment in SGs in cells lacking m^6^A. This observation is consistent with a recent study [27]. And It is likely that YTHDF proteins bind to m^6^A to drive the partition of long mRNAs into SGs.

While PHOTON offers numerous advantages over existing technologies to capture subcellular transcriptomic information, there are opportunities for further optimization. One limitation of the current PHOTON protocol is that only a single set of spatial regions can be selected within each specimen, and the data are inherently aggregated over these ROIs. Although this drawback can partially be mitigated by deconvolution algorithms, future versions of PHOTON may enable spatial barcoding of multiple classes of (and potentially individual) spatial regions. Potential approaches for multiplexed target selection include sequential rounds of photocleavage at different targets with *in situ* ligation of target-specific, barcoded PCR handles taking place between rounds. Another limitation of PHOTON is that during photocleavage, the laser passes through the entire thickness of specimen and can potentially uncage cDNA molecules above and below the focal plane outside of the targeted subcellular compartments. Currently, we navigate this effect by sectioning tissues to approximately single cell thickness and optimizing optical conditions. Both the axial and lateral resolution of PHOTON could be further improved by targeting the RNA molecules with two-photon absorption, enabling fully volumetric photocleavage.

In summary, PHOTON is broadly applicable to spatial transcriptomic analyses of many organisms and cell types. Future use of PHOTON in conjunction with other methods with a similar design principle such as PSS [17] may enable muti-modal profiling (e.g., a combination of the transcriptome, genome, and epigenome) of subcellular compartments to shed light on the mechanisms of gene regulation in a spatially resolved manner.

## Methods

### Cell culture

Cells were maintained in DMEM, high glucose, pyruvate (Thermo Fisher Scientific 11995073), 10% heat inactivated fetal bovine serum (Thermo Fisher Scientific 16140-071), and 1% Pen-Strep (Gibco, 15140-122) at 37°C/5% CO_2_. The day before the PHOTON experiment, cells were washed with 1X dPBS, dissociated with 0.05% Trypsin-EDTA (Thermo Fisher Scientific 25300054), and collected into a new tube. Cell density was measured with the DeNovix CellDrop cell counter. Cells were plated in a Cellvis 96-well glass bottom plate (Cellvis P96-1.5H-N) pre-treated with Matrigel Matrix (Corning 354234) diluted in DMEM at a 1:50 dilution.

To induce SG formation, cells were incubated in DMEM containing 0.2 mM sodium arsenite for 50 minutes at 37°C/5% CO_2_. Mettl3 conditional KO MEF Cells were a gift from Dr. Samie Jaffrey at Weill Medical College of Cornell University. To achieve Mettl3 KO, cells were treated with 500 nM 4-OHT for at least 5 days.

### Tissue preparation

8-week-old female C57BL/6 mice were intraperitoneal injected with 5 IU pregnant mare serum gonadotropin (PMSG, Lee Biosolutions Inc 493102.5) and after 48 hours injected with 5 IU human chorionic gonadotropin (hCG, MilliporeSigma CG10). After euthanasia, ovaries from stimulated mice were dissected and washed with 1X PBS for 3 times. Ovaries were then immersed in 4% PFA for two hours at room temperature. After washing with 1X PBS for 3 times, ovaries were transferred into 10% sucrose solution and rotated overnight in the 4°C cold room, followed by the dehydration with 20% and 30% sucrose solution sequentially. Tissues were embedded in optimal cutting compound (OCT) media after kimwiping excess liquid, frozen on dry ice, and then stored at -80°C for later experiments. The ovarian tissue blocks were cut into sections of 10 μm thickness using Leica CM1950 Cryostat and the sections were transferred into glass-bottom dishes (MatTek P35G-1.5-14-C) treated with 0.1% Poly-L-Lysine (Sigma-Aldrich P8920-100ML).

### PHOTON workflow

Samples are fixed in a glass-bottom dish or plate with 0.4% PFA for 10 minutes at room temperature followed by 3 times of 1X PBS wash. Permeabilization was performed using 0.5% Triton X-100 (Sigma Aldrich, T9284-100ML) supplemented with 0.4 U/ μL RNase inhibitor (NEB, M0314L) for 15 minutes at room temperature followed by 3 times of 1X PBS wash. Samples can then undergo sample-specific immunofluorescent staining to label the ROIs (see Immunostaining below).

*In situ* RT was performed in the RT mix (0.5 μM NPOM-RT, 0.5 μM NPOM-RT-Random, 10 U/μL Maxima Reverse Transcriptase (Thermo Fisher Scientific, EP0741), 0.4 U/ μL RNase inhibitor, 1X RT Buffer, and 1mM dNTPs (Thermo Fisher Scientific, R0194)) on a shaker for 30 minutes at room temperature followed by incubation overnight at 37°C. Samples were protected from light during the RT process.

ROIs were visualized on an inverted Nikon CSU-W1 Yokogawa spinning disk confocal microscope using either a 0.95 NA CFI Apochromat LWD Lambda S 20x water immersion objective lens (for tissue) or a 1.15 NA CFI Apo LWD Lambda S 40x water immersion objective lens (for cells). For GCs in the ovarian tissue, RO1s were selected manually. For nucleoli and SGs, the JOBS and GA3 modules in the NIS-Elements AR software were used to enable real-time segmentation of ROIs using custom scripts. Selected ROIs were scanned for targeted photocleavage using an XY galvo scanning module via the Nikon Ti2-LAPP system (Nikon) coupled with a 50 mW 405 nm laser line. For GCs in the ovarian follicles, photocleavage was performed with 37.5 mW laser power and a dwell time of 300 microseconds. For nucleoli and SGs, 2.5 mW laser power and a dwell time of 100 microseconds were used.

Following photocleavage, samples were digested by incubation in reverse-crosslinking buffer (50 mM Tris pH 8.0, 50 mM NaCl and 0.2% SDS) with 1:50 proteinase K (NEB P8107S) for 30 min at 55 °C in the dark. Nucleic acids were then extracted using the NucleoSpin Gel and PCR Clean-up XS Kit (Takara Bio 740611.250).

Ligation adaptors were generated by annealing 10 μM Splint Oligos with 10 μM P7 Handles in STE Buffer (10mM Tris pH 8.0, 50 mM NaCl, and 1mM EDTA) at a total reaction volume of 50 μL. The annealing was performed by heating the oligo mix at 95 °C for two minutes followed by cool-down to 20 °C at a rate of 1°C per minute.

Ligation was performed by mixing 10 μL of the annealed ligation adaptors with the purified nucleic acids. The ligation reaction was also added with 20 U of RNase inhibitor and 1X T4 DNA Ligase Reaction Buffer to a total reaction volume of 47.5 μL. The ligation mix was then incubated for 30 minutes at room temperature on a shaker in the dark. After that, 2.5 μL of T4 DNA Ligase (NEB M0202L) was added to the mix, bringing the total reaction volume to 50 μL. The ligation mix was then incubated for 30 minutes at room temperature on a shaker in the dark.

To pull down cDNA molecules, 10 μL/sample MyOne Streptavidin C1 Dynabeads (Thermo Fisher Scientific 65001) were placed on a magnetic stand and washed once with 1X B&W-T Buffer (5 mM Tris pH 8.0, 1M NaCl, 0.5mM EDTA, and 0.05% Tween 20) and washed twice with 1X B&W-T Buffer supplemented with 0.4 U/μL RNase Inhibitor. After the washes, each 10 μL of the beads were resuspended in 100 uL of 2X B&T Buffer (10 mM Tris pH 8.0, 2 M NaCl, 1 mM EDTA, and 0.8 U/μL RNase inhibitor).

Following ligation, 50 μL of nuclease-free water was added to the ligation mix, bringing the total volume up to 100 μL. Then, 100 μL of the MyOne Streptavidin C1 Dynabeads resuspended in 2X B&T Buffer was added to each sample. The mixture was rotated on an end-to-end rotator for 60 minutes at room temperature in the dark. After incubation, samples were placed on a magnetic stand. The supernatant was removed, and the beads were washed three times with 1X B&W-T buffer supplemented with 0.4 U/μL RNase inhibitor and washed once with STE buffer supplemented with 0.4 U/μL RNase inhibitor.

After the washes, the MyOne Streptavidin C1 Dynabeads were resuspended in 50 μL of template switch mix for a second round of RT. The template switch mix consisted of 1X RT Buffer, 1 mM dNTPs, 10 U/μL Maxima Reverse Transcriptase, and 2.5 μM template switch oligomer (TSO). Resuspended beads were rotated on an end-to-end rotator for 30 minutes at room temperature in the dark, and then were shaken at 300 rpm for 90 minutes at 42°C. The beads were resuspended by pipetting every 30 minutes during the incubation period. After incubation, beads were washed with STE Buffer for 3 times at the room temperature.

After the STE buffer wash, beads were resuspended in 55 μL PCR Mix (1X Q5 Hot Start High-Fidelity 2X Master Mix (NEB M0494L), 0.45 μM PCR_TSO, and 0.45 μM PCR_P7). The PCR reaction was run on a BioRad T100 Thermal Cycler, which was programmed with an initial denaturation step of 98°C for 30 seconds followed by five cycles of 98°C at 10 seconds, 65°C for 30 seconds, and 72°C for 3 minutes. The final extension was 72°C for 5 minutes.

To determine the number of additional cycles needed for the PCR reaction, after the initial 5 cycles of PCR amplification, samples were placed on a magnetic stand and 2.5 μL of supernatant from each sample was combined with 7.5 μL qPCR master mix (1X SYBR Green (Thermo Fisher Scientific S7563), 0.45 μM PCR_TSO, 0.45 μM PCR_P7, and 1X Q5 Hot Start High-Fidelity 2X Master Mix). qPCR was run on a LightCycler 480 II System (Roche Diagnostics 05015243001) in 384-well plate with the following program: 98°C for 30 seconds; 35 cycles of 98°C at 10 seconds, 65°C at 30 seconds, and 72°C at 60 seconds. 72°C for 5 minutes. The number of qPCR cycles needed to reach ⅓ of the saturated signal was used as the additional PCR cycles needed for each sample.

PCR products from the initial 5-cycle amplification were put back into the thermocycler for additional amplification using the number of cycles determined by the qPCR experiment. After the second round of PCR, samples were placed on a magnetic stand. Supernatant was collected and cleaned with 0.8X AMPure XP Beads (Beckman Coulter A63880). The concentrations of purified PCR productions were determined using the Qubit dsDNA HS Assay (Thermo Fisher Scientific Q32854).

Purified PCR products were subject to sequencing library preparation using the Nextera XT Library Prep Kit (Illumina FC-131-1024) following the manufacturer’s instructions. The resulting libraries were cleaned with 0.7X SPRI Select DNA Beads (Beckman Coulter B23317). The concentration and size distribution of the sequencing libraries were quantified using the Agilent D1000 High Sensitivity Screentape Kit (Agilent 5067-5584) on an Agilent TapeStation 2200 system.

DASH [18] was used to eliminate the rRNA and mitochondrial DNA in the sequencing library. Briefly, crRNA oligos were ordered from IDT and pooled to a stock concentration of 100 μM in IDTE Buffer (10 mM Tris and 0.1 mM EDTA). 100 μM DASH tracrRNA were then mixed in equimolar ratio with the pooled crRNA to a final duplex concentration of 10 μM. The duplex was then heated at 95°C for 5 minutes and allowed to cool to room temperature. The crRNA:tracrRNA complex was then aliquoted and stored at -80°C. To create the ribonucleoprotein (RNP) complex, the 10 μM crRNA:tracrRNA duplexes were combined with EnGen Spy Cas9 NLS (NEB M0646M) in equimolar amounts in 1X PBS to a final concentration of 1 μM. The RNP complex solution was then incubated at room temperature for 5-10 minutes for optimal formation of the RNP complex.

To perform in vitro digestion, each sequencing library was diluted to 10 nM in 20 μL nuclease-free water and mixed with 10 μL of 10X Cas9 Nuclease Reaction Buffer and 70 μL of the Cas9 RNP complex. Reactions were incubated at 37°C for 60 minutes. After incubation, 1 uL of proteinase K was added to each reaction, and the mixture was further incubated at 56°C for 10 minutes to release the DNA substrate from the Cas9 endonuclease.

The post-DASH libraries were cleaned up using 0.8X SPRI Select beads. The concentration and size distribution of the sequencing libraries were quantified using the Agilent D1000 High Sensitivity Screentape Kit (Agilent 5067-5584) on an Agilent TapeStation 2200 system. Libraries were sequenced on either an Illumina NextSeq 2000 or a NovaSeq X Plus.

### Immunostaining

To visualize SGs during the PHOTON workflow, cells were incubated on a shaker for one hour at room temperature with the anti-eIF3eta antibody (Santa Cruz sc-137214) at a 1:200 dilution in 1X PBS supplemented with 0.4 U/μL RNase inhibitor. Cells were then washed three times for 5 minutes each with 1X PBS and incubated on a shaker in the dark for one hour at room temperature with of anti-mouse Alexa 647 secondary antibody (1:200 dilution, Thermo Fisher Scientific A-31571) supplemented with 0.4 U/μL RNase inhibitor. Cells were washed three times for five minutes each in 1X PBS, and post-fixed with 0.4% PFA in the dark for ten minutes at room temperature. After fixation, samples were washed three times for five minutes each with 1X PBS.

GCs in the ovarian follicles can be directly visualized through the fluorescent signals on the RT primers. Nuclei were visualized by staining the tissue with DRAQ5 dye (abcam ab108410) at a 1:300 dilution in 1X PBS for 30-60 minutes at room temperature. Nucleoli were visualized using 2.5 μM nucleolus bright green dye (Dojindo Laboratories c511).

### SNR quantification

Approximately 5,000 HeLa cells per well were used in this experiment. A fixed percentage of cells were photocleaved for each well and the samples were subject to the PHOTON workflow. The amount of the resulting library from each well was quantified and compared with that from the unblocked well (i.e., no photocleavage) using qPCR. For photocleavage, two different laser intensities were assessed. Low laser intensity was set at 2.5 mW with 100 microsecond dwell time. High laser intensity was set at 25 mW with 100 microsecond dwell time.

### HCR-FISH

To validate the HeLa cell SG PHOTON dataset, we used HCR RNA-FISH targeting *AHNAK, MN1*, and *ZACN*. Briefly, sodium arsenite-treated HeLa cells were fixed with 4% PFA for ten minutes at room temperature. After fixation, cells were washed with 1X PBS twice for five minutes each at room temperature on a shaker. Cells were then permeabilized with ice-cold 70% Ethanol overnight at -20°C. After permeabilization, cells were washed twice for five minutes each with 1X PBS and incubated in antibody buffer (Molecular Instruments) for 60 minutes at room temperature on a shaker. Cells were then incubated overnight at 4°C with the anti-eIF3eta antibody diluted at 1:100 in Antibody Buffer.

After primary antibody incubation, cells were washed three times for five minutes each with 1X PBS and incubated for 60 minutes at room temperature with the anti-mouse secondary antibody (Molecular Instruments) diluted at 1:100 in antibody buffer. After incubation, cells were post-fixed with 4% PFA for ten minutes at room temperature and washed twice for five minutes each with 1X PBS, and then washed twice for five minutes each with 5X SSC buffer (Thermo Fisher Scientific AM9770). Cells were then pre-conditioned in pre-warmed hybridization buffer (Molecular Instruments) for 30 minutes at 37°C.

Probes targeting each mRNA were ordered from Molecular Instruments and were diluted to 16 nM in the probe hybridization buffer. These probes were added to the cells for overnight incubation at 37°C. After incubation, cells were washed four times for five minutes each with probe wash buffer (Molecular Instruments) at 37°C and were then washed 5 × 5 minutes with SSCT buffer (5X SSC and 0.1% Tween 20) at room temperature. After washing, cells were pre-conditioned in amplification buffer (Molecular Instruments) for 30 minutes at room temperature.

To amplify the signal, fluorescent hairpins matching each probe set and the antibody were first heated at 95°C for 90 seconds and cooled to room temperature in the dark for 30 minutes. The harpins were then added to the amplification buffer to achieve a final concentration of 60 nM. Cells were incubated with the hairpin solution overnight in the dark at room temperature. After incubation, excess hairpins were removed by washing 5 × 5 minutes with SSCT at room temperature. Cells were imaged on an inverted Nikon CSU-W1 Yokogawa spinning disk confocal microscope with a 1.15 NA CFI Apo LWD Lambda S 40x water immersion objective lens.

### METTL3 inhibitor treatment and m^6^A quantification in HeLa Cells

The METTL3 inhibitor STM2457 (Selleck Chemicals S9870) was used to induce acute depletion of mRNA m^6^A methylation. Cells were treated with 20 μM, 50 μM, and 100 μM STM2457, respectively, while the control cells received the same amount of DMSO. After 6 hours of treatment, cells were washed twice with ice-cold 1X PBS. Total RNA was then isolated using the Direct-zol RNA MiniPrep w/ TriReagent kit (Zymo Research R2053-A). Subsequently, mRNA was isolated from the total RNA using Dynabeads Oligo(dT)25 (Thermo Fisher Scientific #61002). The m^6^A level on mRNAs was then colorimetrically quantified using the m^6^A RNA Methylation Quantification kit (abcam ab185912), with the absorbance signal measured at 450 nm by a CLARIOstar Plus plate reader (BMG Labtech).

### Sequencing data analysis

Raw paired sequencing reads were trimmed to eliminate remaining adapter sequences and bases with quality scores < 25. Sequence with less than 35 bp were also removed. Trimmed data were aligned to the reference genome using HiSAT2 [31]. Features (genes, transcripts and exons) were counted using featureCounts [32]. Pairwise differential expression analysis was performed using DESeq2 [33].

## Data Availability

The raw sequencing data supporting the findings of this study are available in the NCBI BioProject database with BioProject ID PRJNA1152621.

## Code Availability

Custom code is available at https://github.com/HaiqiChenLab/PHOTON.

## Acknowledgments

H.C. acknowledges support from the Cecil H. and Ida Green Center for Reproductive Biology Sciences Endowment and NIH-NHGRI (R01HG013358).

## Author Contributions

H.C. conceived and supervised the project. S.R. and Q.X. performed experiments. Q.C. and X.Z. prepared and processed tissue samples. H.C., F.C., and S.M.M. designed the method prototype. H.C., Q.X., and S.R. analyzed data. H.C., Q.X., and S.R. wrote the manuscript with input from all authors.

## Competing Interests

H.C., S.M.M, and F.C. are on a patent related to this work. F.C. is an academic founder of Curio Biosciences and Doppler Biosciences, and scientific advisor for Amber Bio.

**Figure S1.**
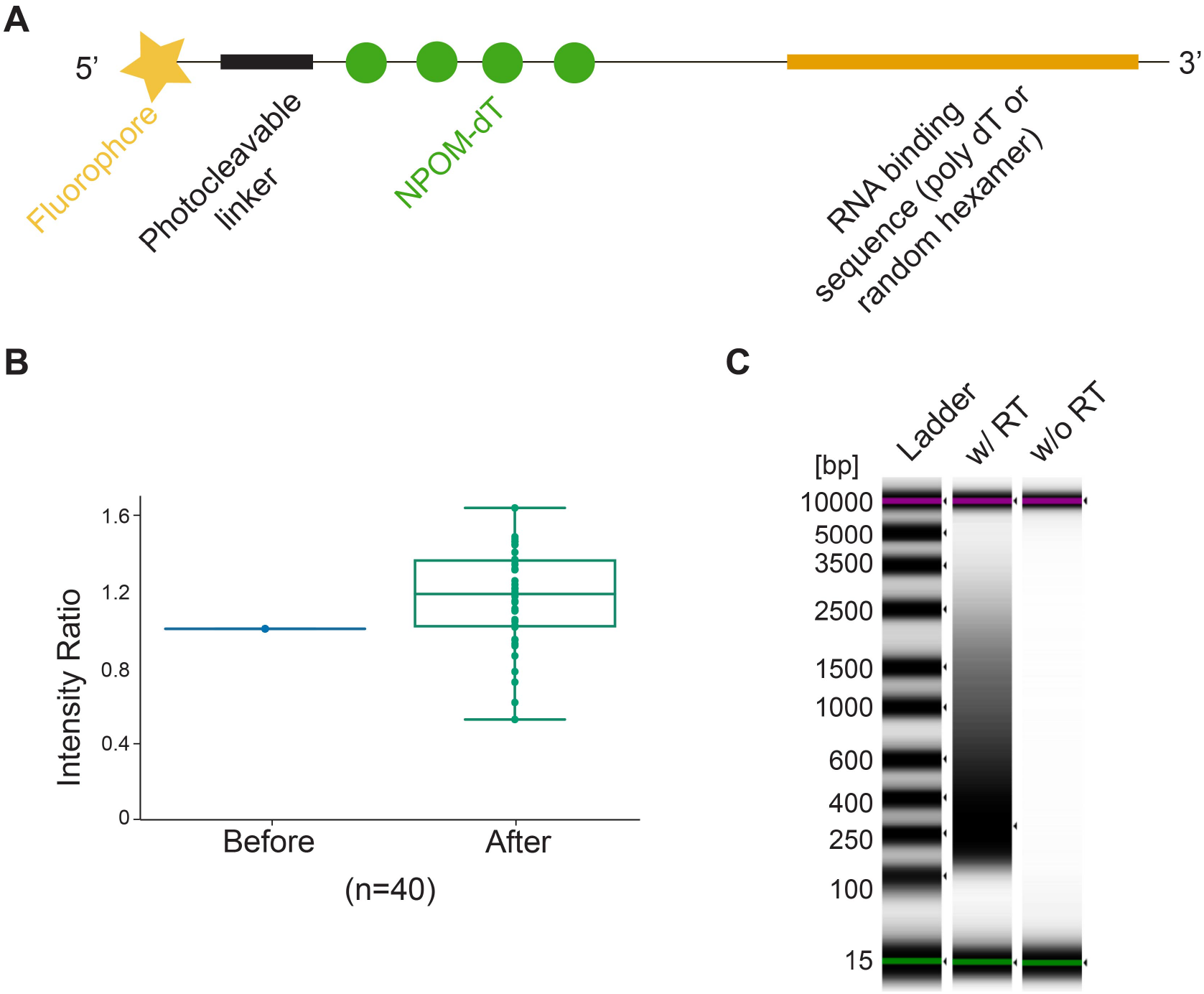
PHOTON RT primer design and the feasibility of PHOTON. **(A)**Design of the RT primer. **(B)**The fluorescence intensities of cells that were not exposed to near-UV laser light remained unchanged after the photocleavage procedure. (n = 40 cells, 1 experiment). **(C)**Tapestation gel image of PHOTON libraries generated with and without RT.

**Figure S2.**
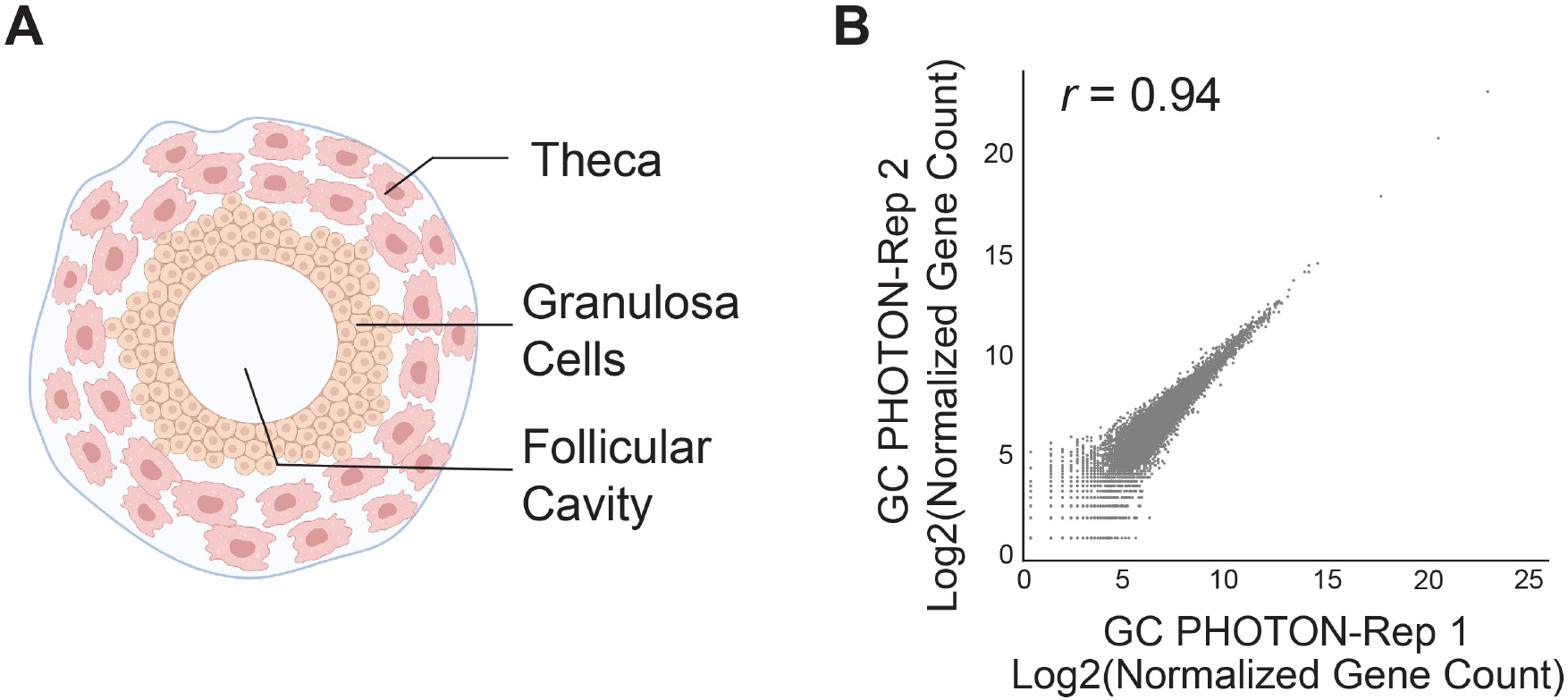
Capturing the GC transcriptome *in situ* using PHOTON. **(A)**Schematic of an ovarian follicle. **(B)**Correlation of the GC transcriptome data between two PHOTON replicates.

**Figure S3.**
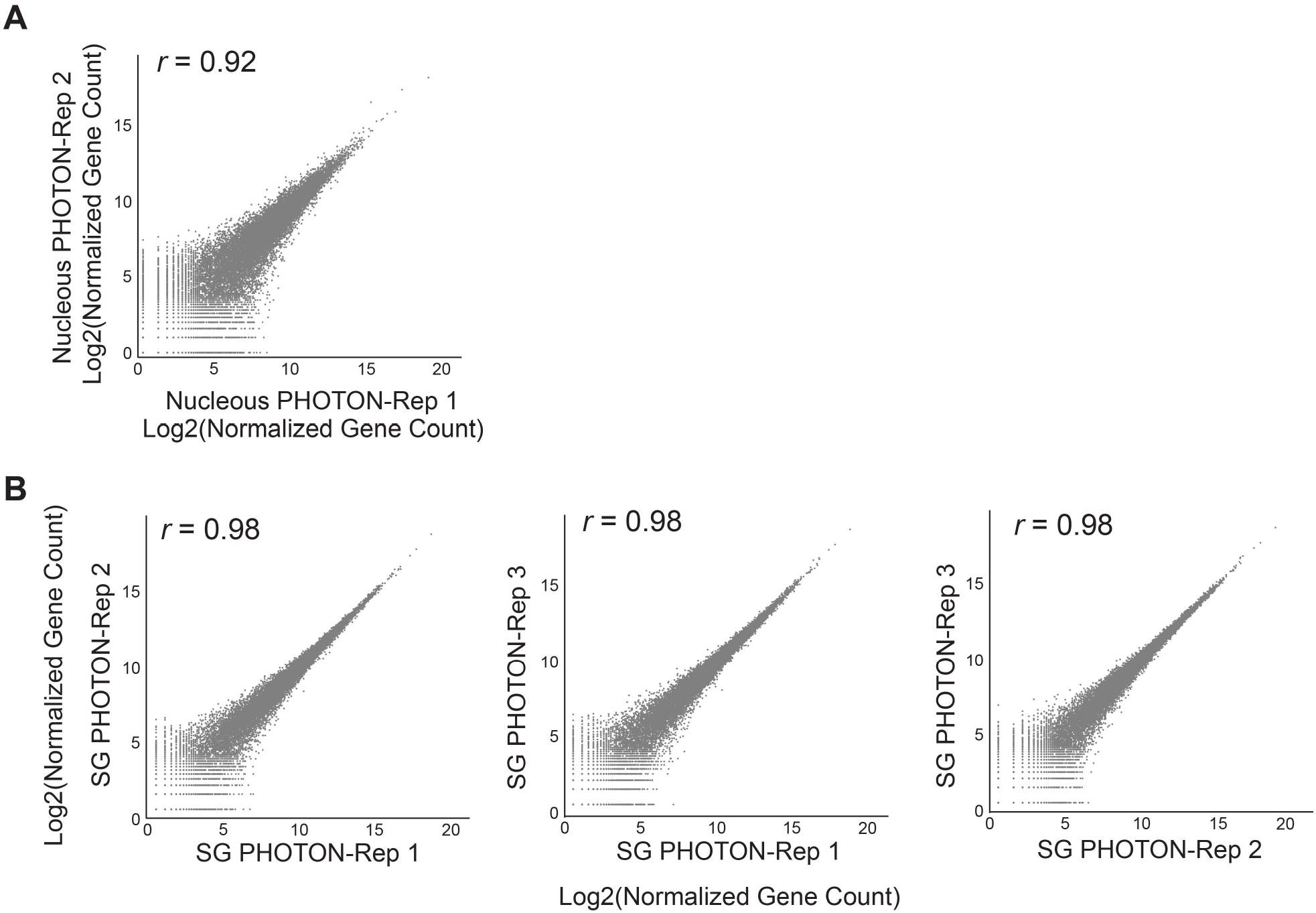
High reproducibility of PHOTON libraries. **(A)**Correlation of the nucleolar RNA content between two PHOTON replicates. **(B)**Correlation of the SG RNA content between three PHOTON replicates.

**Figure S4.**
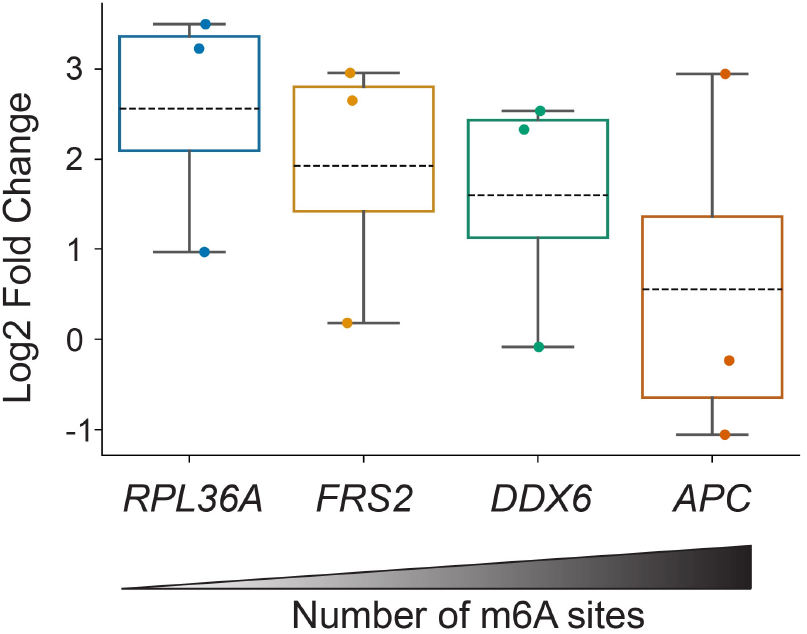
Effects of m^6^A loss on the SG enrichment of selective mRNAs with different number of m^6^A sites. Expression values of four representative transcripts quantified by qPCR were normalized to the ratio of difference in expression from WT and *Mettl3*KO SG relative to that in WT and *Mettl3*KO whole cell mRNA to generate fold change values. The negative effect on SG enrichment in *Mettl3*KO cells was increasingly pronounced in transcripts with more m^6^A.

